# Simultaneous Sensing and Stimulation in Neuromodulation Systems: Quantifying and Mitigating Charge Accumulation Effects

**DOI:** 10.1101/2025.04.30.651467

**Authors:** Heather Orser, Preston Doan

**Affiliations:** University of St Thomas, St Paul, MN 55105 USA; Michaud Cooley Erickson, Minneapolis, MN 55402 USA

**Keywords:** Charge transfer, circuit analysis, low-noise amplifier, neuromodulation

## Abstract

This paper describes the operation of a low-noise amplifier and neurostimulation circuit when both functions are used simultaneously. The design of this circuitry along with the circuit model used to investigate system interactions is described. Expected circuit operation is explored in conjunction with the impact of charge storage at the electrode tissue interface. The test results for the system during independent use of the amplifier and during simultaneous use of the amplifier and stimulator are presented. As seen in the circuit analysis, test results confirm that the charge storage that occurs in the tissue/electrode interface used for stimulation interferes with the measurement of the neural signals of interest when the two circuits share electrodes. By optimizing the configuration used for the amplifier during high frequency stimulation, biological signals can be measured; however, the effective number of bits (ENOB) will be degraded as a function of the stimulation parameters, limiting the applications in which signal amplification can be meaningfully used when sharing an electrode with stimulation circuitry.

## I. INTRODUCTION

Neuromodulation has become a crucial therapeutic approach for a broad range of medical conditions, including chronic pain, Parkinson’s disease, epilepsy, urinary incontinence, and sleep apnea. Traditional neuromodulation systems utilize open-loop stimulation, where fixed parameters like pulse width, frequency, and amplitude are applied without real-time adjustment based on physiological feedback. However, there has been a substantial shift toward closed-loop neuromodulation systems capable of sensing physiological signals and adjusting stimulation parameters in real time to optimize therapeutic outcomes [1]-[3].

In recent years, clinical studies have demonstrated the potential efficacy of closed-loop neuromodulation systems in improving patient outcomes. For instance, closed-loop spinal cord stimulation systems have been shown to provide effective pain relief by dynamically adjusting stimulation parameters based on evoked potentials [1]. Similarly, closed-loop deep brain stimulation (DBS) systems have been used to modulate Parkinsonian symptoms based on beta-band oscillatory activity [2]. Additionally, closed-loop responsive neurostimulator systems for epilepsy have shown promising results in seizure reduction by monitoring real-time neural activity and delivering targeted stimulation [3]. While these systems demonstrate the utility of closed-loop control, they do not utilize shared electrodes for sensing and stimulation, primarily due to the significant technical challenges associated with signal fidelity during concurrent operation.

The challenge of simultaneous sensing and stimulation using a shared electrode has been previously addressed in cardiac pacemakers [4]-[6]. In these devices, a single electrode is commonly used for both pacing and sensing cardiac signals. However, the requirements in pacemakers differ significantly from neuromodulation systems. In pacemakers, pacing frequencies typically do not exceed 5 Hz [7], allowing long intervals between stimulation pulses during which sensing can occur with minimal interference. In contrast, neuromodulation therapies often use stimulation frequencies up to 130 Hz [8], resulting in significantly higher overlap between sensing and stimulation periods. This higher stimulation frequency, combined with smaller signal amplitudes in neural signals, makes the problem of shared electrode sensing in neuromodulation far more complex.

A fundamental challenge in neuromodulation is the large amplitude difference between stimulation and physiological signals. Stimulation pulses often exceed one volt, while neural signals of interest are typically on the order of a millivolt or less. This amplitude disparity introduces substantial signal contamination during concurrent stimulation and sensing. The use of shared electrodes in neuromodulation is further complicated by charge accumulation at the electrode-tissue interface, which can significantly degrade sensing quality. Despite these challenges, sharing electrodes remains highly desirable due to limitations in electrode count and the spatial relevance of sensing sites to the location of stimulation therapy.

Several prior studies have investigated sense and stimulation circuits for neuromodulation applications [9]-[13]. However, all of these circuits use separate electrodes for sensing and stimulation. Furthermore, no previous studies have modeled and experimentally validated the impact of accumulated charge storage at the electrode-tissue interface on sensed signal quality. This work fills this gap by providing a detailed theoretical model, experimental validation, and practical design considerations for simultaneous sensing and stimulation using shared electrodes.

A prior publication by the authors [14] presented the initial design of the custom integrated circuit (IC) used in this study, demonstrating its basic operation for neuromodulation purposes. However, that work did not explore the impact of shared electrodes on sensing quality nor provide detailed design considerations to mitigate charge accumulation effects. This work expands upon the initial IC characterization to specifically address the signal degradation caused by charge buildup at the electrode-tissue interface and to present practical methods for improving sensing performance.

In this paper, we specifically analyze the reduction in the effective number of bits (ENOB) in a sensing amplifier caused by charge storage at the electrode-tissue interface. Using a custom integrated circuit (IC) that supports both sensing and stimulation functions, we characterize how stimulation parameters such as pulse width, frequency, and current amplitude affect the quality of sensed signals. Additionally, we validate these findings using a tissue phantom setup to simulate realistic bio-impedance characteristics. Our results provide a clear quantification of sensing degradation and offer practical design recommendations to mitigate these effects.

The remainder of this paper is organized as follows: Section II describes the circuit design and theoretical model for charge accumulation. Section III presents the test setup and experimental results. Section IV discusses the practical implications of our findings, and Section V concludes the paper with potential future work.

## II. Circuit Design and Analysis

The integrated circuit (IC) described in [14] was fabricated using a 0.18-µm CMOS process for implantable applications, operating from a 3V battery with minimal external components. The IC integrates both stimulation and sensing circuitry, with each configurable to interface with predefined electrodes through multiplexers (MUX). The system architecture, depicted in Fig. 1, consists of a DC-DC converter, stimulation circuitry, sensing circuitry, and four multiplexers that enable electrode selection. External components for stimulation include a charge pump capacitor, C_CAP, and a recharge capacitor, D_CAP. The DC-DC converter employs a single external inductor to generate supply voltages up to 12.5V using a non-inverting buck-boost topology. While the system supports up to eight electrodes plus a reference electrode, this work focuses on configurations where one or two electrodes are shared between the sensing and stimulation circuits.

**FIGURE 1.**
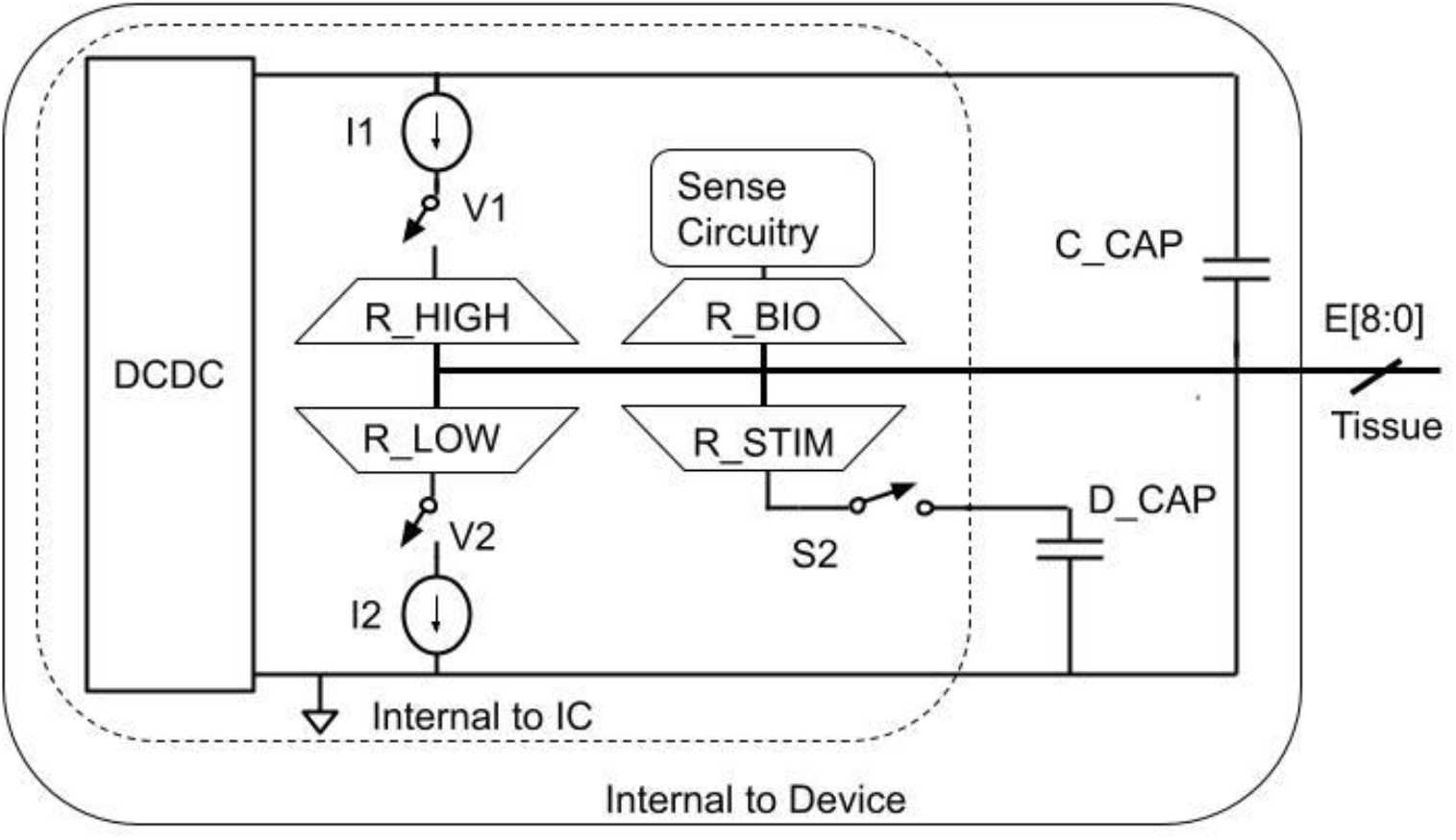
Block diagram of custom IC. External components required for operation of the IC are included for the DCDC Converter, Stimulation Circuitry, and Sense Circuitry. Components required for operation of other blocks have not been included.

### A. Stimulation Engine

The IC generates stimulation waveforms like those in Fig. 2. Each stimulation cycle consists of a stimulation phase, a charge balancing phase, and an inter-pulse interval. During the stimulation phase, charge is delivered to the electrode, followed by a charge balancing phase in which an equal and opposite charge is applied to mitigate electrochemical effects [15]. An inter-pulse interval separates successive pulses to complete the waveform. The stimulation train is configurable in terms of pulse width, amplitude, and frequency, allowing patient-specific therapeutic adjustments. Pulse widths range from 30.518 µs to 976.576 µs, determined by a 32.768 kHz precision clock. Current amplitude is programmable between 0 and 10 mA through a 10-bit digital-to-analog converter (DAC) with 0.22% accuracy and a compliance voltage of up to 12.5V. Stimulation frequency is adjustable based on therapy requirements.

**FIGURE 2.**
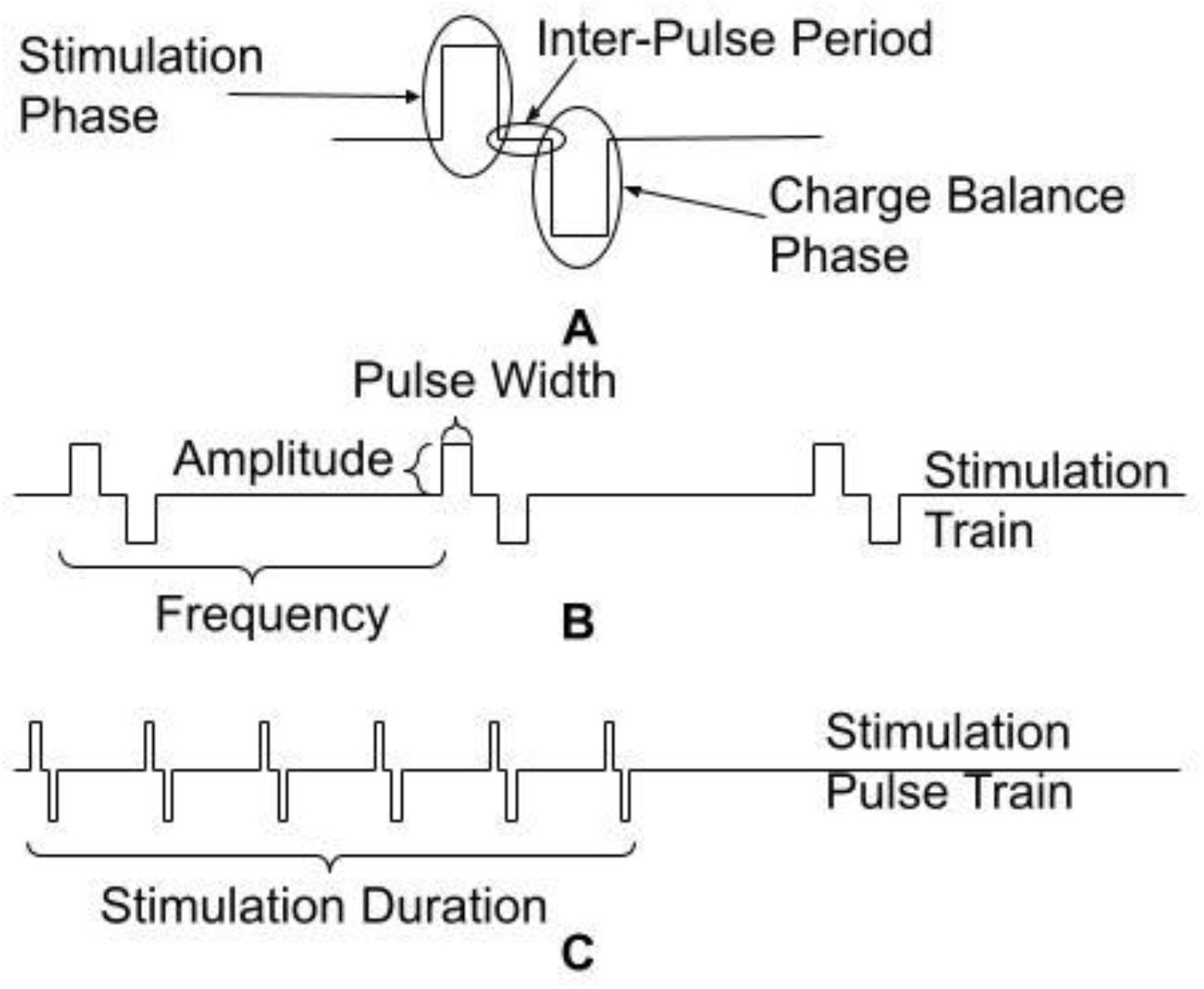
Description of stimulation pulses. Each stimulation pulse consists of a stimulation phase, an inter-pulse period, and a charge balancing phase (A). These pulses with a defined pulse width, amplitude, and frequency are combined into a stimulation train (B). In some cases, this stimulation train is pulsed on and off during therapy and creates a stimulation pulse train (C).

In addition to continuous pulse trains, the stimulator supports burst-mode operation, in which stimulation periods alternate with non-stimulation intervals, a scheme used in neuromodulation therapies such as hypoglossal nerve stimulation [16]. To ensure charge balance at the electrode interface, the IC undergoes functional calibration during manufacturing, where trimmed current sources are stored in non-volatile memory to provide consistent charge balancing.

Electrode selection is managed by the R_HIGH, R_LOW, and R_STIM multiplexers. During the stimulation phase, switches V1 and S2 are closed, allowing current flow through the tissue and accumulation of charge on D_CAP. In the inter-pulse interval, V1 and S2 open to halt current flow. During the recharge phase, S2 and V2 close to discharge D_CAP, restoring charge neutrality at the electrode.

### B. Sensing Amplifier

The sensing amplifier is implemented as a differential amplifier with three programmable gain settings of 100, 1,000, and 10,000. A single-ended representation of the amplifier is shown in Fig. 3. The amplifier has a bandwidth of 0.3 to 500 Hz and supports a programmable sampling rate of up to 1024 samples per second. The input voltage range is determined by the supply voltage and gain setting, supporting input signals up to 30 mV. Electrode selection for sensing is controlled via the R_BIO multiplexer.

**FIGURE 3.**
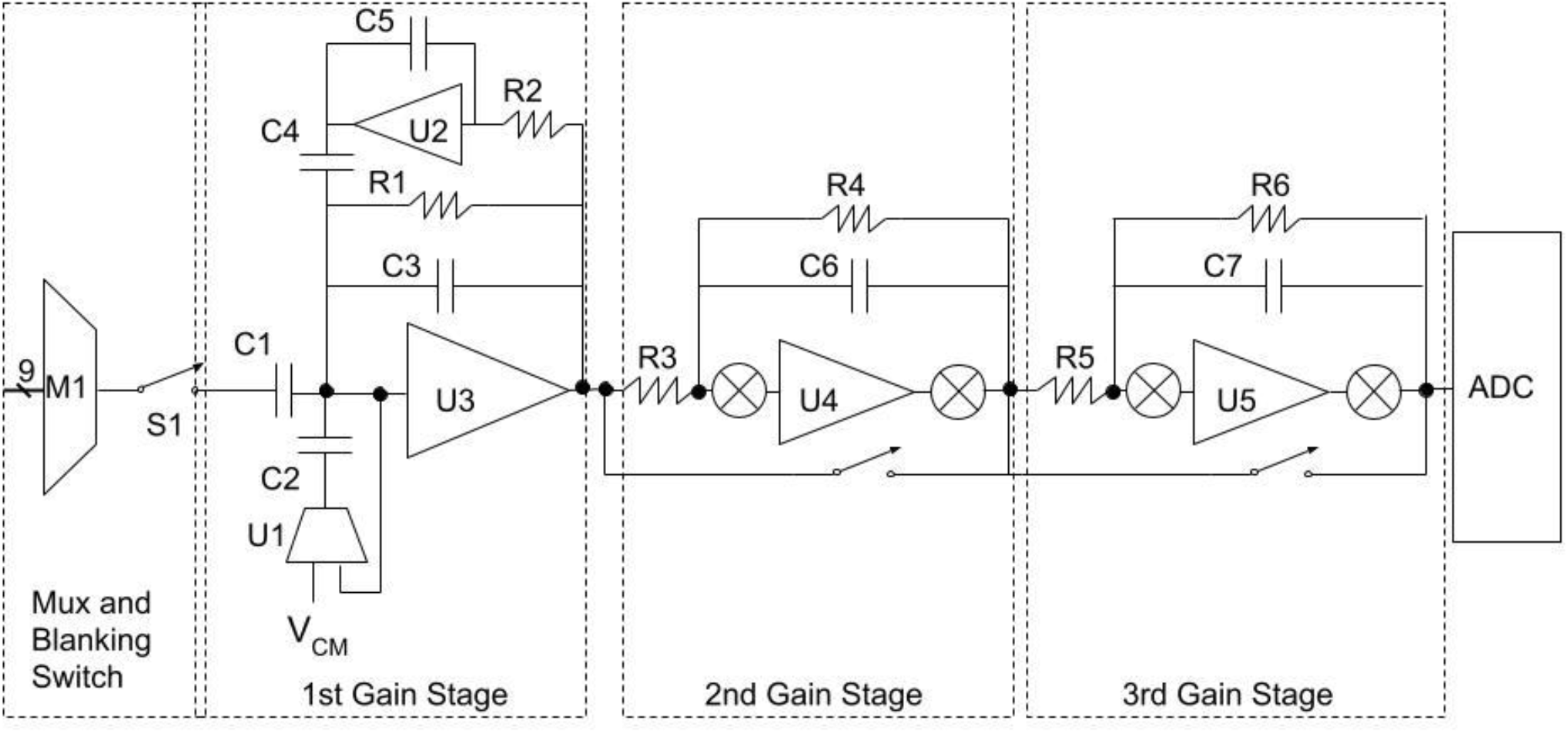
The sense amplifier [12] used for acquiring signals from the body. To reduce the impact of stimulation pulses on amplifier saturation, the input is disconnected from the amplifier for the duration of the stimulation pulse and reconnected at the conclusion. The first gain stage of the amplifier is designed for large common mode disturbances to further account for normal operation during stimulation while allowing for low noise operation appropriate to capture cardiac and neural signals. Subsequent stages allow additional amplification and offset regulation as appropriate for the application.

To mitigate interference from stimulation artifacts, the amplifier integrates a blanking switch that disconnects the inputs during stimulation pulses. This switch, designed to tolerate stimulation pulses up to 12.5V, opens prior to stimulation onset and remains open throughout the pulse duration, thereby isolating the amplifier input from high-voltage stimulation transients. The blanking period is programmable, allowing extended isolation if necessary. Once the predefined isolation period ends, the switch closes, restoring connectivity to the electrodes.

The amplifier consists of three cascaded gain stages. The first stage provides a gain of 100 and incorporates a common-mode feedback amplifier that allows the system to tolerate common-mode perturbations of up to 100 mV. The second and third stages each provide a gain of 10 and can be bypassed to configure the overall system gain. For characterization, the default gain setting is 100, as this stage dominates the input-referred noise in all configurations. Under these conditions, the system is capable of processing signals with amplitudes up to 30 mV, ensuring high-fidelity acquisition for bioelectrical signal analysis.

### C. Analog to Digital Converter (ADC)

Once the signal is amplified, it is quantized by a 12-bit ADC at rates up to 1024 samples/sec. Sampling can be performed as a multiple of the stimulation rate or at frequencies asynchronous with the stimulation rate. Asynchronous sampling samples the signal at the output of the amplifier at a rate independent of the stimulation frequency. Synchronous sampling can sample the signal up to four times per stimulation pulse at a defined time after the stimulation pulse. This configuration prevents sampling during a stimulation pulse.

Regardless of the type of sampling used, the blanking switch opens the connection between the electrode and the input of the amplifier during the stimulation phase. For synchronous sensing the amplified signal is guaranteed not to sample during a stimulation pulse. With asynchronous sensing ADC sampling will collide with some stimulation pulses. This situation is illustrated in Fig. 4.

**FIGURE 4.**
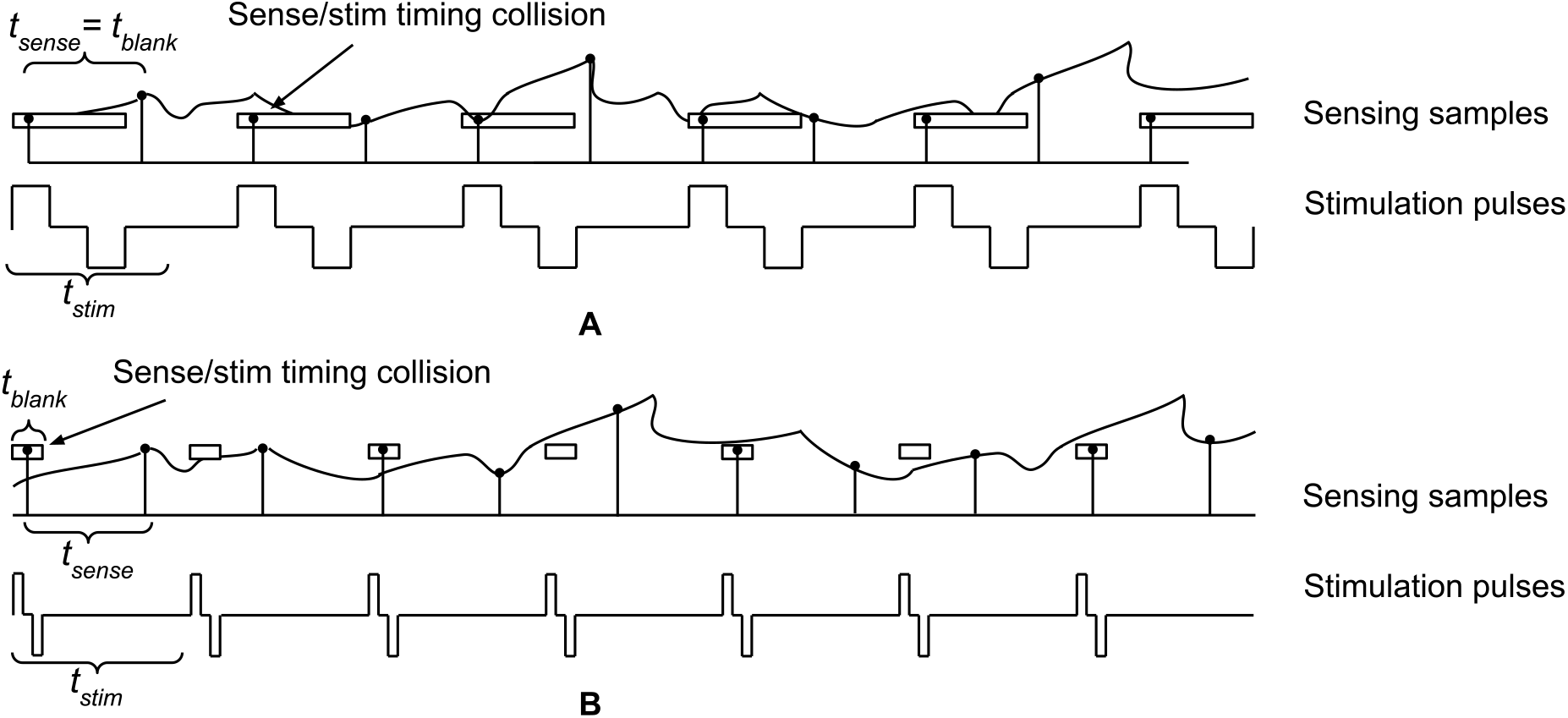
The timing diagram of sampling time for the sense amplifier and stimulation pulse timing. In the top diagram, the period of a stim pulse is 2 times the period of a blanking duration and sensing time is equal to blanking time. As can be seen, this results in 1 sample data for every 2 samples, or 1/2 of samples are data. The bottom diagram shows a period of the stim pulse equal to 6 blanking periods and the sense rate equal to 4 blanking periods. In this case there are 2 samples taken for every 3 samples, or 2/3 of samples are data.

To calculate the percent of samples that occur during a stimulation period, the following assumptions are made: sampling occurs instantly at a rate of 1/*tsense*, the blanking switch is open for time *t_blank_*during a stimulation pulse train, and the stimulation period is *t_stim_*. If *t_blank_*is equal to *t_sense_*, as illustrated in Fig. 4, the percent of samples taken that occur when stimulation is off can be calculated as:

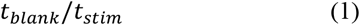

When *t_blank_*is less than *t_sense_*, it will take multiple stimulation periods before a sense sample occurs during stimulation. For instance, if the blanking duration is 250 µs, the sampling rate is 1 ms, and the stimulation period is 1.5 ms, the percent of samples taken that are data can be calculated in terms of *t_sense_*and *t_stim_*as:

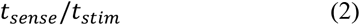

This is the general form of the expression and addresses situations where the stimulation pulse is greater than the sense sampling period. Fig. 4 illustrates the timing associated with each of these timing situations. Collisions that occur during sampling cause harmonics of the stimulation frequency in the sensed signal, reducing the spur-free dynamic range (SFDR) of the sampled signal. Due to this, it is expected that sampling the signal synchronous with the stimulation pulses will result in improved SFDR.

### D. Sense and Stimulation Configuration

The sense and stimulation circuitry described previously can be configured to support sharing one or more electrodes for both functions. A simplified circuit diagram with previously described sense and stimulation circuits sharing at least one electrode can be drawn as shown in Fig. 5A and 5B. The circuit diagram includes an electrical model of the electrode/tissue connection that consists of resistors and capacitors and has been previously described [15]. The electrode/tissue model represents the ionic interface present at an electrode and can be built with each electrode model meeting at a common point. The capacitance, denoted C_dlx_, and the faradic resistance, denoted Z_fx_, are properties of the interface between the electrode and the ionic solution while the solution resistance, denoted R_s_, is a property of the solution surrounding the electrodes. Typical values of these components for a chronic implant are C_dlx_ between 1 µF – 200 µF, Z_fx_ of approximately 1 MΩ, and R_s_ between 25 – 4000 Ω. Exact values of each element depend on the surface area of the interface and the electrode and electrolyte materials. Fig. 5A shows the connections when the sense amplifier and the stimulation circuitry share only one electrode and Fig. 5B shows the connections when the sense amplifier and the stimulation circuitry share both electrodes.

**FIGURE 5.**
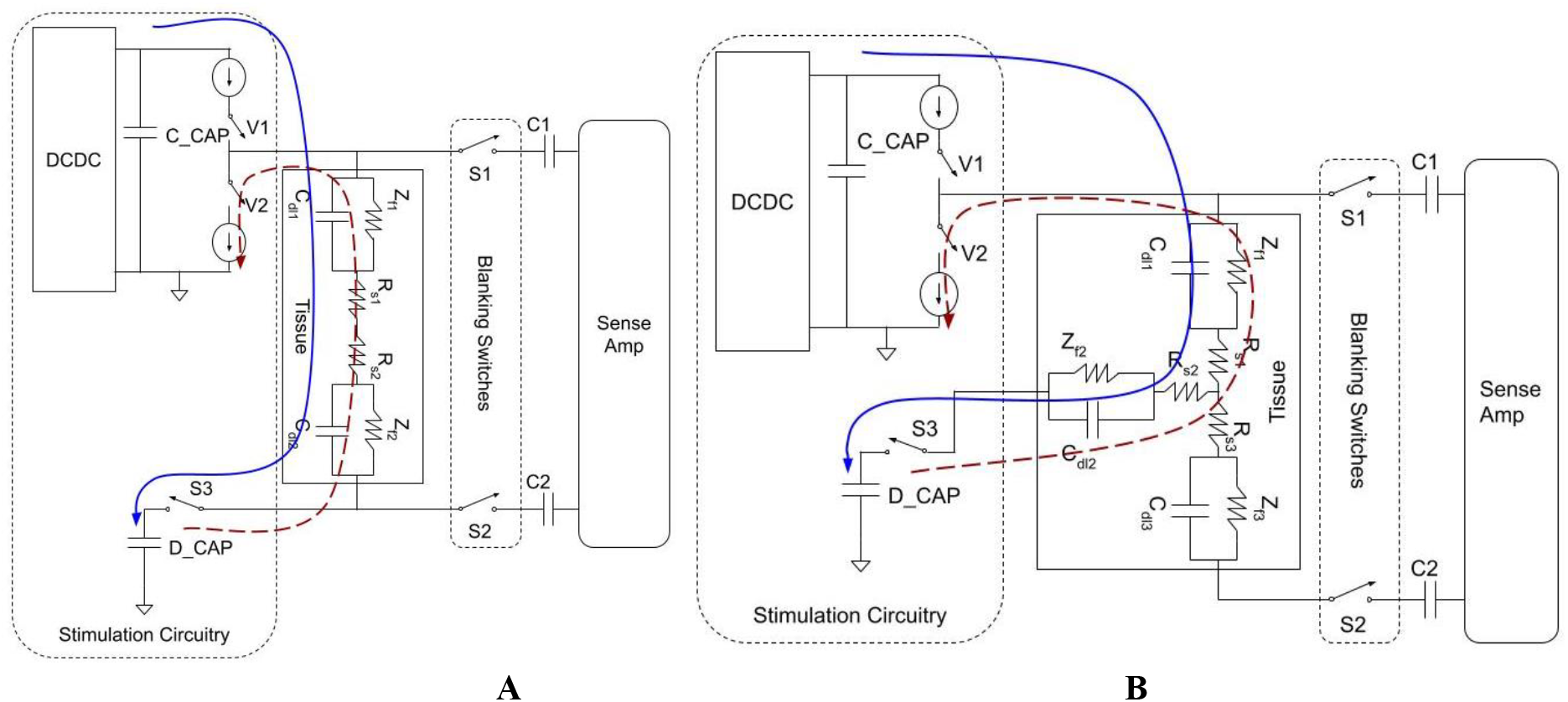
Circuit configuration for sensing when both electrodes are shared (A) and when one electrode is shared (B). Solid line shows path of cathodic current and dashed line shows path of anoodic current during a stimulation pulse. Switch timing for the circuitry is V1 and S3 close during cathodic current delivery to tissue, then V1 and S3 open for the interpulse interval, followed by V2 and S3 closing for the anodic pulse. Once the stimulation pulse is complete, S1 and S2 close to enable sensing.

The circuits shown in both of these configurations operate as follows. During delivery of the stimulation phase, switches V2, S1, and S2 are open and switches V1 and S3 are closed, causing charge transfer from capacitor C_CAP through the tissue to capacitor D_CAP. At the completion of the stimulation period, all switches are opened for the duration of the inter-pulse period. For the recharge phase, switches S3 and V2 are closed and charge is transferred from capacitor D_CAP back to the circuit ground. At the conclusion of the recharge phase, all switches are open for a programmable hold off period and then switches S1 and S2 on the sense amplifier are closed to allow sensing of neural based signals to resume.

In both the Fig. 5A and 5B configurations, some charge is accumulated in charge storage elements at the electrode/tissue interface. The charge storage elements, C_dlx_, for commercially available clinical electrodes, typically have values of 1-3 µF. Each element has a discharge path through Z_f_; however, with a value of approximately 1 MΩ, the discharge time constant is roughly 1 second, much longer than the period of a neural stimulation signal. The result of this is that stimulation pulses can produce an offset voltage build up on capacitors C_dlx_ which is present at the input to the sense amplifier.

The offset voltage on C_dlx_ capacitors is illustrated in Fig. 6 with an offset voltage V_stim_offx_ shown across the respective capacitor. This offset voltage can be calculated by evaluating the stimulation and charge balance phases of a stimulation pulse. In Fig. 6A and 6B, during the stimulation phase current moves from electrode E2 to E1, inducing a voltage change on the capacitor *C_dlx_*equal to

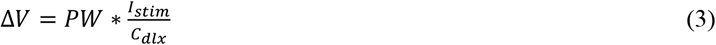

**FIGURE 6.**
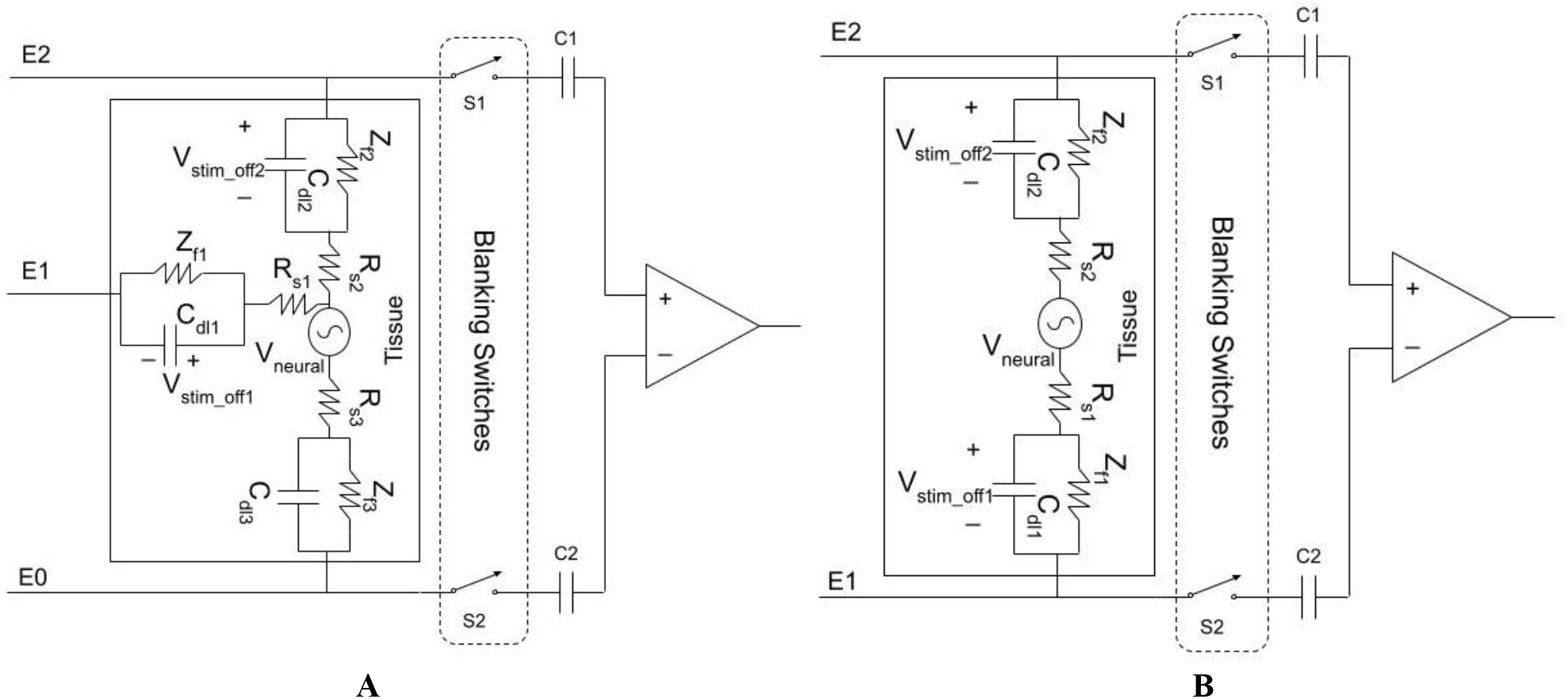
Impact on sensing of offset voltages induced on electrodes by stimulation when both electrodes are shared (A) and when one electrode is shared (B). V_stim_offx_is shown on capacitors C_dl1_and C_dl2_in the case where stimulation is applied between E1 and E2. For sensing performed between E1 and E2 (A), the input voltage to the amplifier is now V_neural_+ V_stim_off1_+ V_stim_off2_. Similarly, in (B) where sensing is performed between E0 and E2, the input voltage to the amplifier is V_neural_+ V_stim_off2_,. Both of these configurations effectively reduce the useable dynamic range of the amplifier with a larger effect when both electrodes are shared.

where *I_stim_*is the current through the tissue, *DV* is the voltage change across the capacitor, and *PW* is the pulse width of the stimulation pulse. The charge balance portion of a stimulation pulse reverses the current through the tissue. For a perfectly balanced pulse the voltage change on *C_dlx_*is equal and opposite to the voltage during the stimulation pulse, resetting the voltage on the capacitor to zero. However, mismatch between the stimulation and charge balance phases results in a finite offset voltage that is stored on the electrode, resulting in a gradual increase in voltage on C_dlx_ with each stimulation pulse. As stimulation signals repeat, this results in a growing offset voltage, denoted V_stim_off1_ and V_stim_off2_, across each electrode used for stimulation. For typical neuromodulation stimulation frequencies, charge builds up on *C_dl1_*and *C_dl2_*to a value of approximately

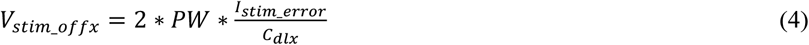

where *I_stim_error_*is the error in amps between source and sink currents. This voltage is now seen at the input to the sense amplifier for any electrodes shared between the sense and stimulation circuitry. For the purpose of safety this charge build-up is a concern if the accumulated charge is sufficient to induce irreversible Faradaic reactions [15] and consequently some system designs to limit the impact of charge build-up have been proposed including using capacitors to ensure charge balance and electrode shorting [17]. While these solutions can significantly reduce or eliminate the apparent charge at the stimulation interface, any charge that remains on *C_dlx_*after application of these solutions will result in an offset voltage that impacts the sensed signal.

For the sense amplifier pictured in Fig. 6A, there is a voltage at the input of *V_neural_+ V_stim_off2_*, The presence of the offset voltage on the capacitors sets the least significant bit (LSB) size for the system and results in a maximum effective number of bits (ENOB) equal to

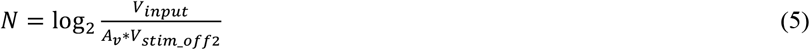

where *A_v_*is the gain of the signal before the ADC and *V_input_*is the input voltage range of the ADC. The result of this charging is that the resolvable range of the sense amplifier will be reduced due to the offset voltage *V_stim_offx_*. If both electrodes are shared with the sensing circuitry, as see in Fig. 6B, this will further reduce the sensing resolution due to the presence of both *V_stim_off1_*and *V_stim_off2_*at the input of the amplifier. In this case, the ENOB of the system amplifier is reduced to

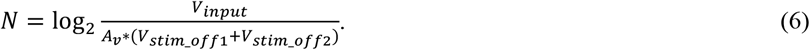

Overall, using the same electrode for both sense and stimulation functions is expected to result in reduced signal sensitivity that is related to the charge storage at the electrode/tissue interface and the imperfections in the stimulation circuitry matching. These two features result in a voltage offset at the input of a sense amplifier which effectively limits the minimum observable signal.

## III. Test Results

The base performance of the stimulator and sense amplifier of the circuit were previously assessed and described in [14]. The purpose of the assessment below is to characterize the operation of the system relative to expected performance when one or two electrodes are shared between sense and stimulation functions.

Six sense and stimulation settings were initially assessed in a benchtop set-up using a resistive load with two shared electrodes, as shown in Fig. 7A. A factorial test was performed to identify any settings that exacerbate reductions in performance. Based on the settings tested with a resistive load, settings were chosen for testing with a tissue phantom, as shown in Fig. 7B and C. These results are compared to the theoretical impact of charge storage on ENOB.

**FIGURE 7.**
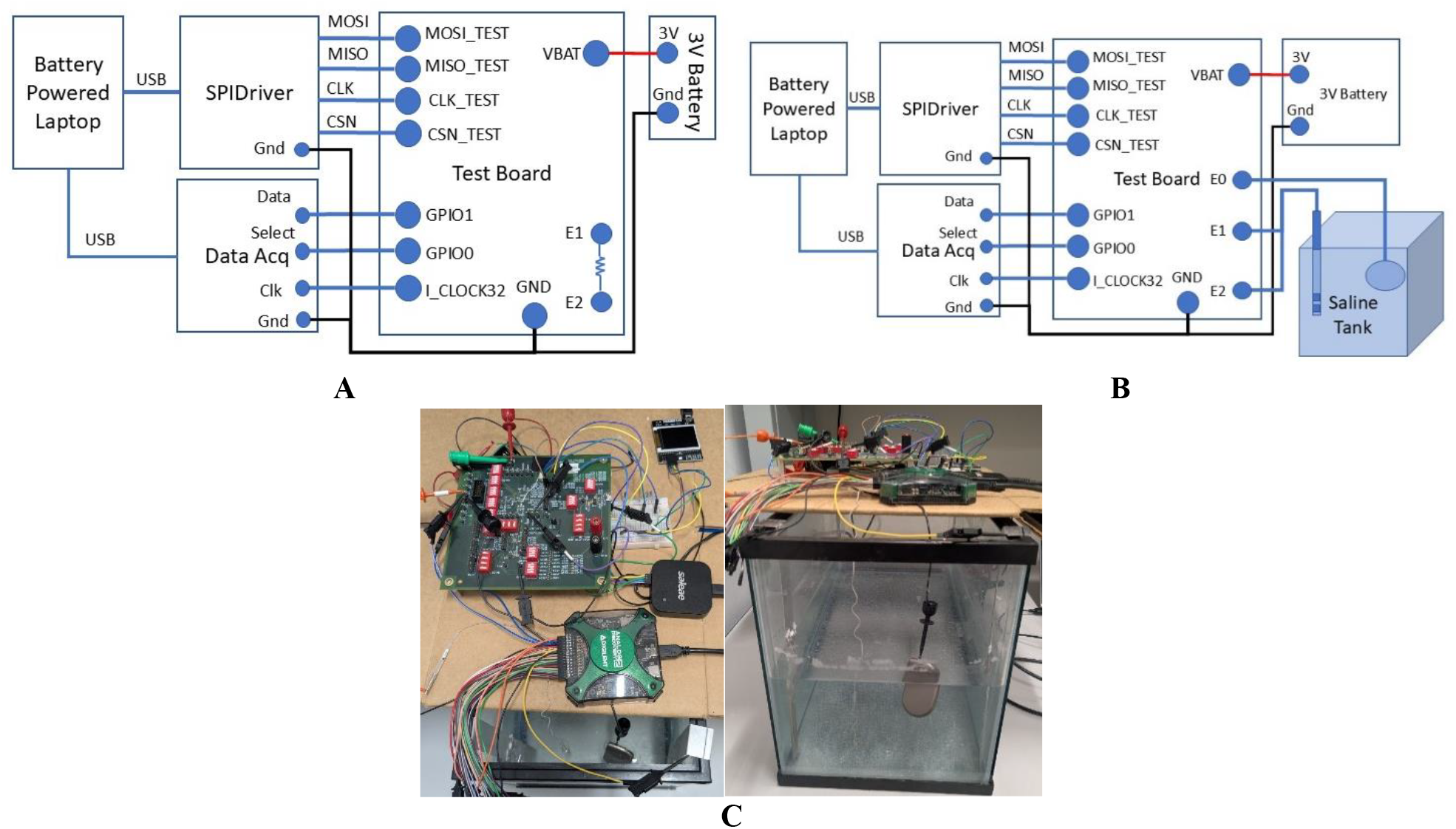
Test configurations used for assessment of sense and stimulation interaction. All equipment is powered by battery to remove 60 Hz impact on system operation. Commands for configuring the sense and stimulation system are delivered via a serial peripheral interface (SPI) and the digital data out of the sense amplifier is collected using a digital analyzer. Stimulation signals were initially assessed with sense and stimulation delivered using the same electrodes through a resistor (A). Based on the results of initial testing, a subset of configurations are tested using the configuration shown in (B) where stimulation is delivered to the electrodes on a lead in a tank filled with saline and the sense amplifier is configured to amplify the signal between one of the lead electrodes and an indifferent electrode in the tank. The test setup used for tank testing is shown in (C).

### A. Factorial Test Design and Results

A factorial test was designed to assess sense and stimulator settings in the test configuration seen in Fig. 7A. This test configuration uses a 1 kΩ resistor between electrodes E1 and E2. The stimulation circuitry drives a signal across this load and the sense circuitry monitors the signal between these electrodes. No excitatory signal apart from the stimulation pulse is driven across the load, so if the sense circuit is well decoupled from the stimulation circuitry during the sampling period the observed signal will be a noise floor. Due to the absence of charge storage in the load, this test is designed to assess the contribution of factors such as cross-talk to a reduction in dynamic range.

For this configuration, a factorial test to assess six variables was used to characterize the output of the sense amplifier during the delivery of stimulation. The range of values used are representative of ranges used in neuromodulation therapies and are described in Table I.

**TABLE I.**
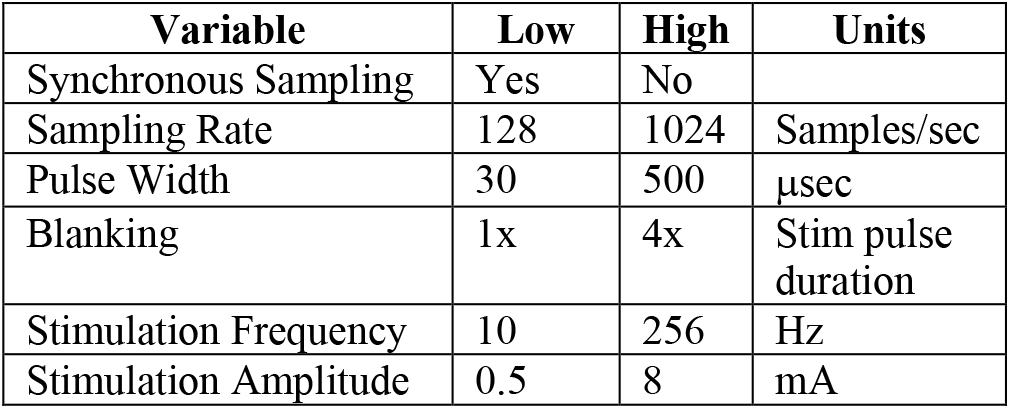
Variables assessed in test configuration shown in Fig. 5A.

The frequency spectrum of all test cases was calculated to assess the impact of stimulation on the sense signal. Two representative frequency plots of the signal output are shown in Fig. 8A and 8B. In the first figure, the sense amplifier is sampling synchronously with the stimulation pulse at a rate of 1024 Samples/sec while stimulation occurs at 128 Hz with an 8 mA, 30 µs stimulation pulse. For this configuration no stimulation pulses are observed across the frequency spectrum. In Fig. 8B the sense amplifier is sensing asynchronously with the stimulation pulse at a rate of 1024 Samples/sec while stimulation occurs at a frequency of 100 Hz with an 8 mA, 30 µs stimulation pulse. This configuration clearly illustrates stimulation pulse intrusion on the sense signal. Both signals have the same pulse width, blanking period, stimulation amplitude, and sampling rate. The only difference between these signals is the stimulation frequency. This example is illustrative of the impact of sampling the sense data during a stimulation pulse.

**FIGURE 8.**
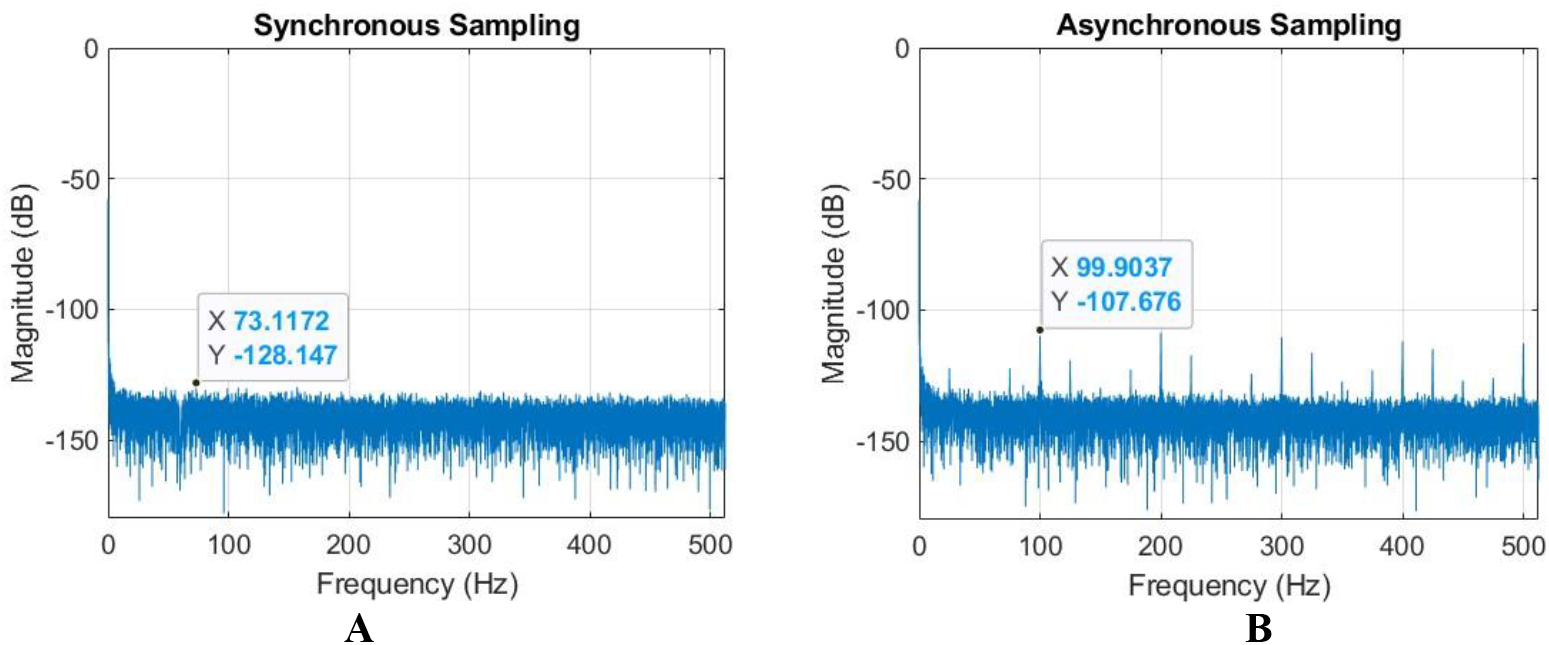
Representative frequency plots of sense output for testing using the configuration shown in Fig. 6B. The spectrum in A is taken using a sampling time that is synchronous with stimulation pulses while the spectrum in B uses asynchronous timing

For each factorial test configuration the spur free dynamic range (SFDR) was calculated and used to analyze the factors that impact sensing behavior. Some reduction in SFDR was observed with higher frequency stimulation and significant reduction in sense performance occurredd when using asynchronous sampling. Other tested factors had a non-significant impact on overall sensing performance. These outcomes along with required therapy settings were used to define system configurations for testing with a tissue phantom.

### A. Phantom Test Design and Results

The system was tested in a saline phantom injected with a sinusoidal signal to assess the behavior of the system with simultaneous sensing and stimulation. Settings representative of therapeutic settings were used for the stimulation pulse while sensing settings were chosen as synchronous with the stimulation pulse with a 1x blanking period. The signal was sampled four times for each stimulation pulse resulting in a sampling rate of 1024 Samples/sec. Testing was performed with a 50 Hz signal injected into the saline tank using four configurations: sensing 50 Hz between E0 and E1 with and without stimulation between E1 and E2 and sensing 50 Hz between E0 and E1 with and without stimulation between E0 and E1. A subset of the results can be seen in Fig. 9.

**FIGURE 9.**
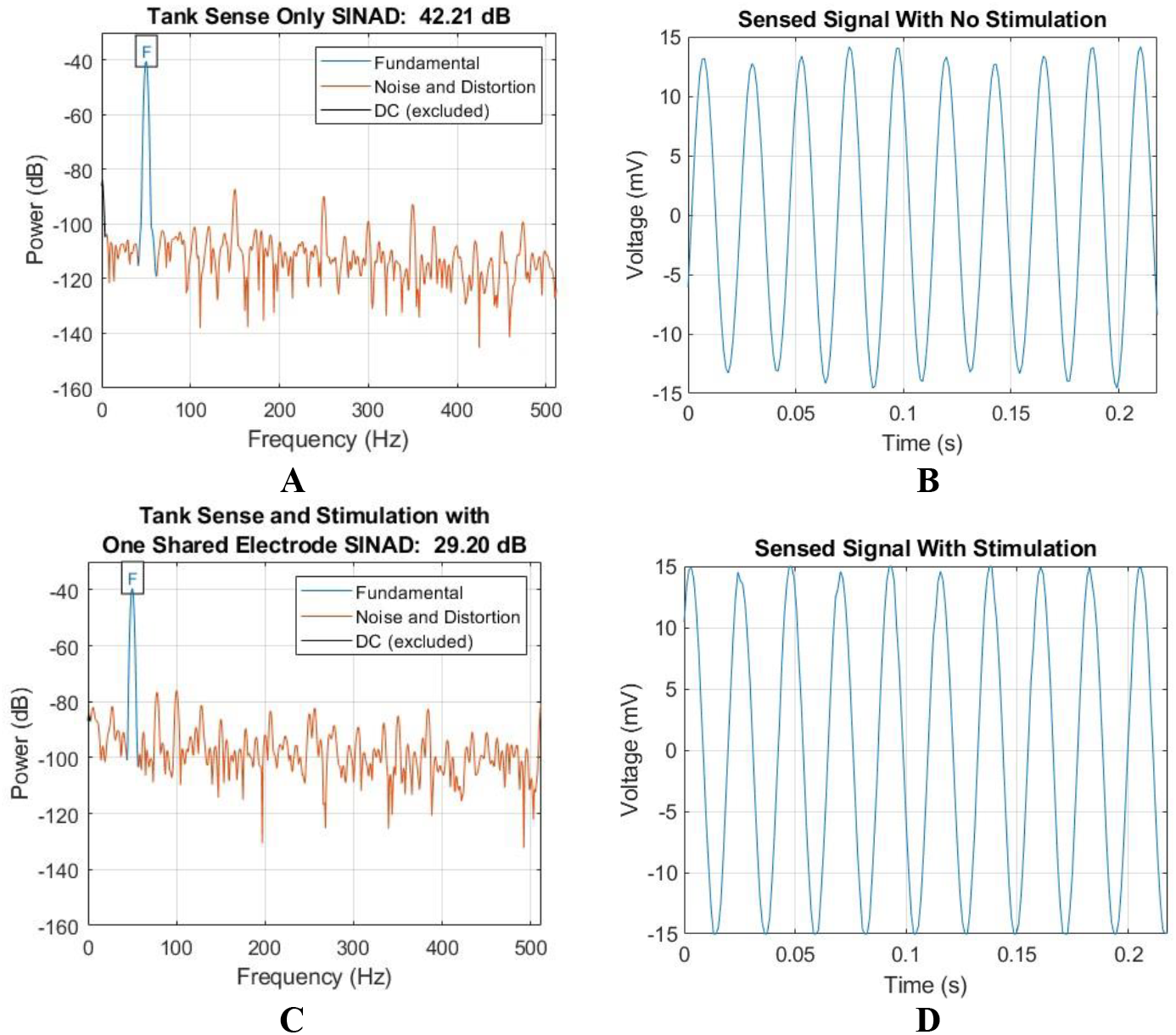
Representative SINAD and time domain plots of sense output for testing using the configuration shown in Fig. 8B. The signal in A and the time domain image in B are taken between the case and an electrode to quantify the injected signal quality. The plots shown in C and D are measurements of the signal between the case and an electrode when stimulation is present between the electrodes on the lead. Note that while the time domain signal appears sinusoidal, stimulation artifact can be seen in the frequency domain.

The values of *C_dlx_*and *Z_fx_*in the test system were characterized and found to be approximately 1.3 µF and 1 MΩ respectively for E1. Source and sink current outputs from the stimulator are calibrated to be within 0.22%. Using (4) and the stimulation parameters to be applied, the expected resolution of the system can be predicted. For a stimulation pulse of amplitude 0.5 mA, pulse width 30 µsec, and frequency 128 Hz produced between E1 and E2 which shares one electrode with the sense amplifier, as shown in Fig. 6A, the expected V_stim_off1_ seen by the amplifier would be approximately 25.4 µV. The time constant to dissipate the charge is calculated using the measured values of *C_dl_*and *Z_f_*and found to be 1.3 seconds, much longer than the stimulation period. As a result, when stimulation pulses are applied at a frequency of 128 Hz charge accumulation is expected on *C_dl1_*during each pulse. This effectively reduces the number of bits that can be resolved in the system.

For the sense amplifier operating from a 3 V supply with a gain of 100, the number of resolvable bits can be calculated by (5) as 10.2, a reduction of 1.8 bits from the system design of 12.

Test results with this configuration can be seen in Fig. 9. For both configurations a 50Hz signal was injected in the tank using aluminum rods to emulate a neural signal. The frequency response of the signal in a sensing only test can be seen in Fig. 9A and the time domain signal is shown in Fig. 9B. The DAC used to generate the 50 Hz signal contained higher order harmonics and the effective circuit formed by the rods with the saline acts as a high pass filter, resulting in a signal to noise and distortion ratio (SINAD) of 42.21 dB. This represents the upper bound of system performance for this test.

Fig. 9C shows the frequency response of the system when stimulation is applied. The SINAD is observed to be 29.2 dB, a reduction of 13.01 dB from the sense only assessment in Fig. 9A with clear stimulation intrusions in the frequency spectrum. This SFDR change effectively reduces the ENOB of the system by 2.15-bits, slightly higher than the predicted impact of charge storage on the electrode.

For the test configuration where E0 and E1 are used for both sensing and stimulation, the expected V_stim_off1_ as seen in Fig. 7B remains approximately 25.4 µV; however, this configuration also has a V_stim_off2_. The electrode used for E0 in this configuration is a much larger electrode representing a reference electrode in an implant. This larger electrode has *C_dlx_*and *Z_fx_*of approximately 20 µF and 1 MΩ respectively. For these values, the value of V_stim_off2_ is calculated as 3.3 µV. Using (6), this means the ENOB of the system is predicted to be 10-bits, slightly less than the ENOB for the imbalanced system. From test results, the SINAD difference between the two configurations was 16 dB, or 2.66-bits, a slightly bigger signal loss than the 2-bit reduction predicted.

## IV. Discussion

### A. System Operation

Using the same electrode for both sense and stimulation functions is desirable for many applications due to the simultaneous need both to manipulate the neural circuit in the vicinity of a given electrode and to monitor the behavior of the neural circuit. As described in this paper, the behavior of charge at the electrode interface presents a challenge to this goal. By carefully assessing the timing and functionality of the sense circuitry and understanding the stimulation parameters an approximate impact on the effective dynamic range can be calculated.

For the system described and tested using typical therapeutic levels of stimulation, signals can be measured with a resolution of 25.4 µV during ongoing stimulation when there is one shared electrode and a resolution of 28.7 µV when there are two shared electrodes. These signal levels may provide meaningful measurement capability for signals such as evoked compound action potentials (ECAP) and electro-cardiograms (ECG) where signal amplitudes can reach up to 8 mV [18] and 5 mV [19], respectively. At maximum amplitudes this would result in a meaningful signal to noise ratio (SNR) that can be useful for applications such as measuring heart rate or determining ECAP timing. These signals could be used to adjust therapy levels in applications where the neural response provides an indication of therapy effectiveness or they could provide additional health information for a patient [20].

Additionally, from (4) we can see that improving mismatch between source and sink current supplies can improve the system performance. Because mismatches have no direct impact on therapy, improving matching between source and sink offers the best option for improving SNR when sharing electrodes with sense and stimulation.

The voltage offset can be further improved by adjusting the signal current or pulse width; however, current and pulse width parameters are critical for effective therapy. Consequently, this is unlikely to be a preferred option due to the impact on therapeutic settings.

Finally, a system that employs shorting between the stimulation electrodes could significantly reduce effective stored charge on an electrode. Doing this requires additional switches and requires a brief period of time before resuming sensing but could result in significant benefit when both sense and stimulation functions share electrodes by discharging the stored charge at the tissue interface.

### B. System Limitations

When it is necessary to use the same electrode for both sense and stimulation, there is a clear limitation in system operation. If the designed system limits the attainable sensed signal, diagnostic use cases may require that measurements be made with stimulation turned off or that measurements be made in a configuration where no electrodes are shared between the sense and stimulation circuitry.

## V. Conclusion

In this work, we have presented a detailed analysis of the impact of shared electrodes on sensing performance in neuromodulation systems, specifically quantifying the reduction in effective number of bits (ENOB) due to charge accumulation at the electrode-tissue interface. By characterizing the relationship between stimulation parameters, charge accumulation, and offset voltage, we demonstrated that increasing pulse width and stimulation mismatch exacerbates the degradation of sensing accuracy. Our results show that without sufficient charge cancelation, the resulting DC offset significantly impairs signal resolution, reducing the amplifier’s dynamic range. Practical mitigation strategies such as improving current matching and using synchronous sampling can improve performance, but fundamental limitations remain. Future work will explore advanced hardware solutions, such as electrode shorting to further enhance the sensing capability on for electrodes shared with stimulation. This study provides critical insight into the design trade-offs for shared-electrode neuromodulation systems and offers a foundation for developing more effective closed-loop therapeutic devices.

---

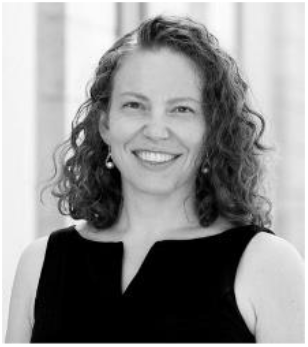

**Heather Orser** received her BSEE from Minnesota State University, Mankato in 1999 and her MSEE and PhD from the University of Minnesota. She is currently an assistant professor of Electrical and Computer Engineering at the University of St Thomas, St. Paul, MN.

Prior to her time at St Thomas, Heather worked in the development of implantable neuromodulation systems at both Inspire Medical and Medtronic where she led the development of a number of next-generation systems and successfully assessed the safety of implantable devices for patients undergoing MRIs. At the University of St Thomas, she teaches Circuit Analysis, Introduction to Biomedical Design, Introduction to Engineering, and Senior Design. Her research focuses on the development of neuromodulation systems for use in research and the clinic.

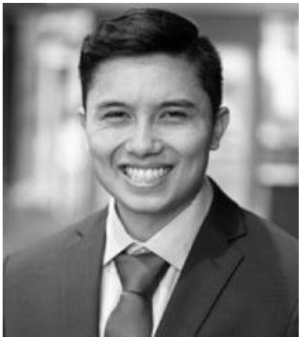

**Preston Doan** received a bachelor’s of science degree in electrical engineering and a bachelor’s of arts in Physics with a minor in Theology from the University of Saint Thomas, Saint Paul, Minnesota in May 2024.

In the summer of 2022 to May of 2023, he worked with Doctor Orser on bio-medical research involving projects like neuromodulation and low-voltage sense and stim devices. The projects worked with the University of Minnesota and Inspire Medical Devices respectively. Some other notable projects he worked on in academia involved lab work in physics like setting up and using optical tweezers and matter-antimatter particle annihilation testing. Outside of academia he worked as a resident advisor for freshmen engineers in which he actively participated in the student life around campus. As of June 17th 2024, he is presently working at Michaud Cooley Erickson (MCE) as an electrical designer in Minneapolis, Minnesota, and is actively working towards getting his PE license.

